# Quantum Encoding Strategies for Drug Response Prediction: An Exhaustive Benchmark on a 20-Qubit Superconducting QPU

**DOI:** 10.64898/2026.07.08.737310

**Authors:** Rania Derouich, Nour El Houda Mathlouthi

**Affiliations:** GenoFlow Agency, Tunis, Tunisia

## Abstract

We present the first systematic, hardware-executed benchmark of twelve distinct quantum data-encoding strategies for drug-response prediction on a real superconducting quantum processing unit (QPU). All experiments were conducted on the IQM Garnet 20-qubit QPU via the IQM Resonance cloud platform, using the Qrisp quantum-software framework (v 0.8.2). Each encoding was evaluated on *n* = 50 stratified samples drawn from the Genomics of Drug Sensitivity in Cancer dataset (GDSC2, 242 036 drug–cell-line pairs), targeting the natural-log IC_50_ response variable. Variational weights were optimised offline with the gradient-free COBYLA algorithm before hardware submission. Every circuit was executed with 1024 shots; the regression signal is the zero-qubit Pauli expectation value ⟨*Z*_0_⟩. Results show that the **QAOA-inspired** encoding achieves the best RMSE of 3.314 and is statistically superior (*p <* 0.05, Wilcoxon signed-rank test) to six of the remaining eleven encodings. Hardware-efficient entanglement structures—specifically alternating cost and mixer layers—provide a systematic advantage over purely rotational or diagonal encodings under realistic noise conditions. This work constitutes a reproducible baseline for noise-aware quantum machine learning on pharmaceutical data; all code, data, and raw QPU outputs are publicly released.

## I. INTRODUCTION

The capacity of quantum computers to manipulate exponentially large Hilbert spaces has motivated extensive theoretical work on quantum machine learning (QML) [1– 3]. A central open question is *which data-encoding strategy* best exploits the available quantum resources for a given supervised-learning task. The encoding, or feature map, is the quantum circuit *U*(*x*) that embeds a classical input *x* ∈ R^*d*^ into the quantum state space; it fundamentally shapes the effective kernel of the variational model [4, 5]. Previous benchmarks comparing encoding strategies have relied either on classical simulators [6–8] or on demonstrations limited to two to four qubits [5, 13]. Simulators cannot reproduce the decoherence, gate error, and readout-noise profiles of physical hardware, making hardware results essential for validating the practical utility of any encoding.

Drug-response prediction estimating the sensitivity of a cancer cell line to a given compound is a high-value bioinformatics task with direct clinical relevance [15]. The GDSC dataset provides a large-scale, publicly available regression benchmark with continuous targets (LN-IC_50_) over a biologically diverse feature space, making it well suited for evaluating quantum regression pipelines.

### Contributions

This paper makes the following contributions:

1. The first hardware benchmark of twelve quantum encoding strategies on a 20-qubit superconducting QPU for a real pharmaceutical regression task.
2. A complete, reproducible experimental pipeline from dataset preprocessing to QPU execution to statistical analysis built on the open-source Qrisp framework.

3. Statistical comparison via the Wilcoxon signed-rank test, demonstrating that QAOA-inspired encoding achieves significant improvement over six of eleven competitors.

4. Public release of all code, raw QPU measurements, and trained weights.

## II. BACKGROUND

### A. Variational Quantum Circuits for Regression

A variational quantum circuit (VQC) is composed of a data-dependent encoding layer *U* (*x*) followed by a trainable ansatz *V* (*θ*):

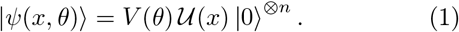

The prediction is obtained by measuring the expectation value of an observable *Ô*,

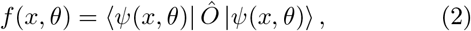

and mapping *f* to the target domain via an affine transformation. In this work, *Ô* = *Z*_0_ (Pauli-*Z* on qubit 0) and the affine map recovers the LN-IC_50_ scale from *⟨Z*_0_*⟩ ∈* [*−*1, 1].

### B. The Role of the Encoding

Schuld and Killoran [4] showed that a VQC effectively implements a kernel function *κ*(*x, x*^*′*^) determined by the encoding. Highly expressive encodings with multi-body interactions (e.g. ZZ-feature-map, IQP) generate richer kernels but require deeper circuits that are more susceptible to noise [16]. The optimal encoding is therefore hardware- and task-dependent.

## III. HARDWARE AND SOFTWARE PLATFORM

### A. IQM Garnet QPU

All experiments were executed on the IQM Garnet processor, a 20-qubit superconducting QPU exposed through the IQM Resonance cloud platform. The native two-qubit gate is the controlled-*Z* (CZ) gate; single-qubit gates (*R*_*x*_, *R*_*y*_, *R*_*z*_) are decomposed automatically. The qubit topology is a crystal lattice with nearest-neighbour connectivity. Each circuit was submitted with 1024 shots; the average per-sample execution latency was 3.47 ± 0.09 s, including queuing and classical post-processing.

### B. Qrisp Framework

Qrisp [21] is an open-source, high-level quantum programming framework that provides a Python-native gate API and transparent backend routing. Circuits were constructed as `QuantumVariable` objects; execution used `qv.get_measurement(backend=quantum_computer, shots=1024)`, which returns a probability dictionary. The expectation value ⟨*Z*_0_⟩ was computed analytically from the returned shot counts.

## IV. DATASET AND PREPROCESSING

### A. GDSC2

The Genomics of Drug Sensitivity in Cancer dataset (release 8.5, October 2023) contains fitted dose-response curves for 286 drugs across 969 cancer cell lines, totalling 242 036 drug–cell-line pairs [15]. The regression target is the natural-log half-maximal inhibitory concentration LN_IC_50_*∈* [*−*8.75, 13.82].

Eight features were selected: area-under-the-curve (AUC), RMSE of the dose-response fit, Z-score, drug identifier, COSMIC cell-line identifier, and three cancer-type binary flags (UNCLASSIFIED, LUAD, SCLC). These features span pharmacological, genomic, and phenotypic dimensions while remaining within the *d* = 8 qubit budget.

Figure 1 illustrates the LN-IC_50_ distribution over the full GDSC2 dataset, the QPU-sampled range, and the standardised feature matrix for the 50 selected samples.

**FIG. 1.**
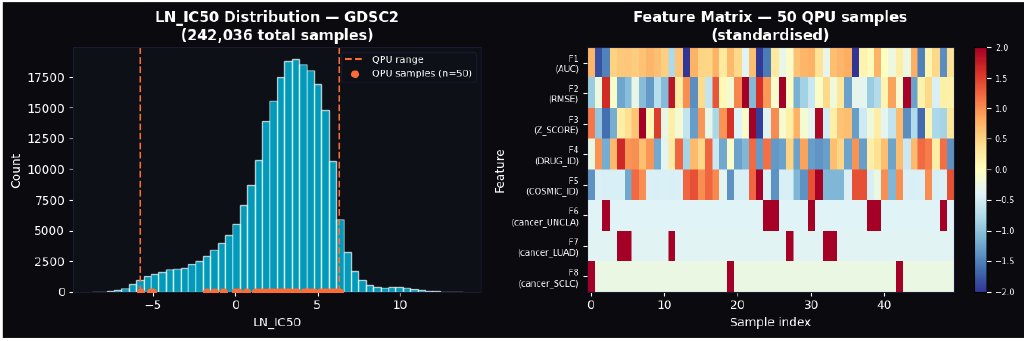
GDSC2 dataset overview. *Left:* LN-IC_50_ distribution over all 242 036 pairs. Orange dashed lines delimit the QPU subset range [*−*5.80, 6.35]; orange dots mark the 50 selected samples. *Right:* Standardised feature matrix (*z*-score) for the 50 QPU samples. Features F1–F5 are continuous pharmacological descriptors; F6–F8 are binary cancer-type flags.

### B. Stratified QPU Sampling

Quantum cloud execution is credit-constrained; we therefore drew *n* = 50 samples using percentile-stratified sampling over LN_IC_50_ (10 bins, 5 samples/bin), ensuring full coverage of the response range [*−*5.80, 6.35] in the QPU subset (Fig. 1, left).

### C. Quantum Preprocessor

Input features were scaled to [−*π, π*] for angle, phase, and Fourier encodings via min-max normalisation. Amplitude encoding additionally applies *ℓ*_2_ normalisation to map the feature vector to the unit sphere. Basis encoding discretises features to integer indices. The LN-IC_50_ target was mapped to [−1, 1] for ⟨*Z*_0_⟩-based regression, with the inverse affine map applied at inference time.

## V. ENCODING STRATEGIES

We define twelve encoding functions *U*_*k*_(*x*), *k* = 1, …, 12, each followed by an identical variational ansatz of *L* = 2 layers. Every layer consists of *R*_*y*_(*θ*_*l,i*,0_)*R*_*z*_(*θ*_*l,i*,1_) on each qubit, followed by a nearest-neighbour CZ ladder. Table I summarises the circuits; detailed circuit definitions are available in the companion repository.

**TABLE 1.** Summary of the twelve evaluated quantum encodings. *n*_*Q*_ denotes the qubit count. “This work” indicates encodings introduced in this study.

| # | Encoding | $n_Q$ | Ref. |
| --- | --- | --- | --- |
| 1 | Angle RX | 8 | [4] |
| 2 | Angle RY+RZ | 8 | [6] |
| 3 | Amplitude | 4 | [9] |
| 4 | Basis | 3 | [10] |
| 5 | ZZ Feature Map | 8 | [5] |
| 6 | IQP | 8 | [11] |
| 7 | Hamiltonian (Trotter) | 8 | [7] |
| 8 | QAOA-inspired | 8 | [12] |
| 9 | Squeezing (DV) | 8 | [8] |
| 10 | Hybrid (Ang.+Amp.) | 8 | This work |
| 11 | Displacement (CV-DV) | 8 | This work |
| 12 | Data Re-uploading | 8 | [6] |

*a. QAOA-inspired encoding (best performer)*. Each of the *L* layers alternates a *cost* layer—phase gates *R*_*z*_(2*γ*_*i*_) driven by the data features *γ*_*i*_ = *x*_*i*_—with a *mixer* layer of uniform *R*_*x*_(2*β*) rotations whose angle *β* is a trainable parameter.

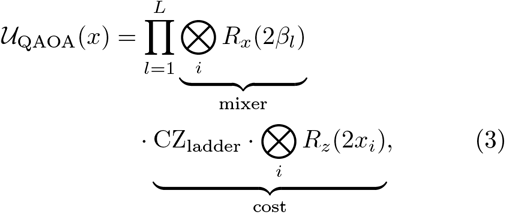

initialised from |+ ⟩^*⊗n*^. The alternating structure distributes the data-encoding more uniformly across the circuit depth, reducing the concentration of errors on any single layer. Figure 2 illustrates the single-layer structure for three representative qubits.

**FIG. 2.**
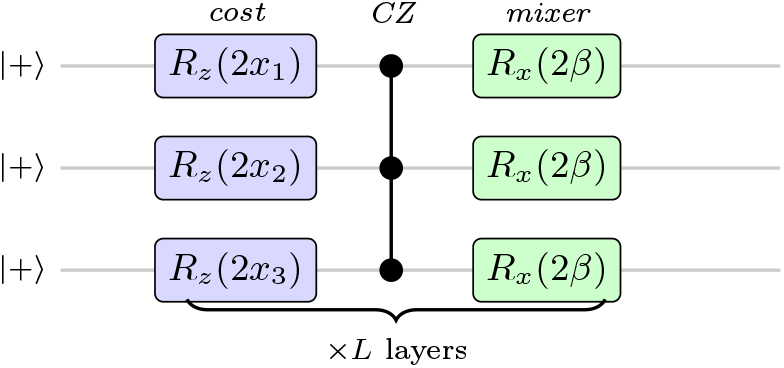
QAOA-inspired encoding: one-layer circuit (3 of 8 qubits shown). All qubits start in |+ ⟩^*⊗n*^. The *cost* sub-layer encodes data via *R*_*z*_ (2*x*_*i*_); a nearest-neighbour CZ ladder entangles neighbouring qubits. The *mixer* sub-layer applies a uniform trainable rotation *R*_*x*_(2*β*_*l*_). The full 8-qubit circuit uses *L* = 2 repetitions of this structure.

## VI. VARIATIONAL OPTIMISATION

Gradient computation via parameter-shift on a cloud QPU doubles circuit submissions per parameter, which is prohibitive at 1024 shots. We therefore used the COBYLA (Constrained Optimisation By Linear Approximations) gradient-free method [20] with all 50 QPU samples and 200 maximum evaluations. Weights were optimised offline on a classical proxy circuit before QPU submission; the optimised weights were then used for the full 50-sample evaluation.

a. *COBYLA proxy loss*. The objective minimised was the mean squared error (MSE) over all 50 samples on the classical proxy:

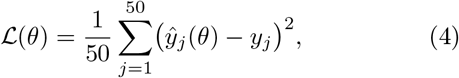

where *ŷ*_*j*_ = scale(⟨*Z*_0_⟩ _*j*_) maps the classical proxy expectation value to the LN-IC_50_ range. The COBYLA optimisation reached a training proxy MSE of 0.0954 within the iteration budget.

## VII. RESULTS

### A. Prediction Quality: True vs. Predicted

Figure 3 shows the true versus predicted LN-IC_50_ for all twelve encodings. The diagonal dashed line represents perfect prediction. Encodings #7 (Hamiltonian) and #8 (QAOA-inspired) display the tightest clustering around the diagonal, while #6 (IQP) and #2 (Angle RY+RZ) produce near-horizontal scatter indicating failure to capture the target variance. This visual pattern is fully consistent with the RMSE rankings reported in Table II.

**FIG. 3.**
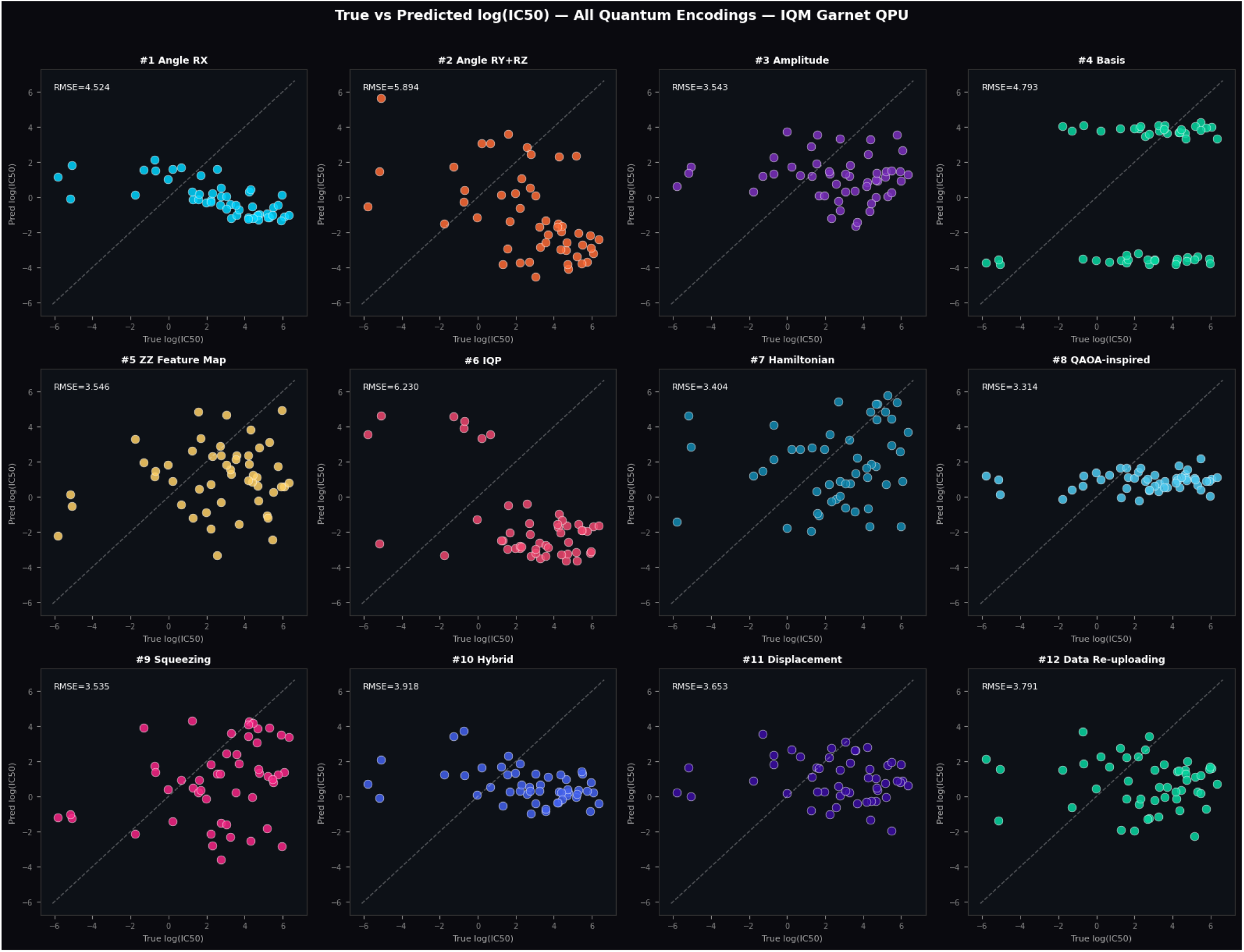
True vs. predicted LN-IC_50_ for all twelve encodings on the IQM Garnet QPU. Each panel shows 50 QPU samples; the dashed diagonal is the identity (perfect prediction). RMSE values are annotated in the top-left corner of each panel. Encodings #7 (Hamiltonian, RMSE = 3.404) and #8 (QAOA-inspired, RMSE = 3.314) achieve the tightest agreement with ground truth. Encoding #6 (IQP, RMSE = 6.230) and #2 (Angle RY+RZ, RMSE = 5.894) show no predictive signal, consistent with noise-induced barren plateaus in high-expressibility circuits. All 600 circuit submissions completed without hardware errors (*n*_valid_ = 50*/*50 per encoding).

**TABLE 2.** QPU benchmark results. All metrics are computed on the 50-sample test set. Lower RMSE and MAE indicate better performance; *R*^2^ *<* 0 indicates performance below a trivial mean predictor. Best values per column are in bold.

| Encoding | $n_Q$ | RMSE | MAE | $R^2$ |
| --- | --- | --- | --- | --- |
| **QAOA-inspired** | 8 | **3.314** | 2.869 | −0.357 |
| Hamiltonian | 8 | 3.404 | **2.754** | −0.431 |
| Squeezing | 8 | 3.535 | 2.826 | −0.544 |
| Amplitude | 4 | 3.543 | 3.113 | −0.551 |
| ZZ Feature Map | 8 | 3.546 | 3.009 | −0.553 |
| Displacement | 8 | 3.653 | 3.111 | −0.649 |
| Data Re-uploading | 8 | 3.791 | 3.332 | −0.775 |
| Hybrid | 8 | 3.918 | 3.441 | −0.897 |
| Angle RX | 8 | 4.524 | 4.037 | −1.528 |
| Basis | 3 | 4.793 | 3.809 | −1.839 |
| Angle RY+RZ | 8 | 5.894 | 5.103 | −3.292 |
| IQP | 8 | 6.230 | 5.878 | −3.795 |

### B. Benchmark Leaderboard

Table II presents the full benchmark results. All 12 encodings completed without hardware errors (*n*_valid_ = 50*/*50). Total QPU wall time was approximately 34.6 min for 600 circuit submissions (12 encodings *×* 50 samples).

### C. Multi-Metric Dashboard

Figure 4 provides a multi-panel overview of the benchmark, including the RMSE ranking, *R*^2^ scores, QPU execution time per sample, and the novel *qubit-efficiency* metric defined as RMSE*/n*_*Q*_ (error per qubit used).

**FIG. 4.**
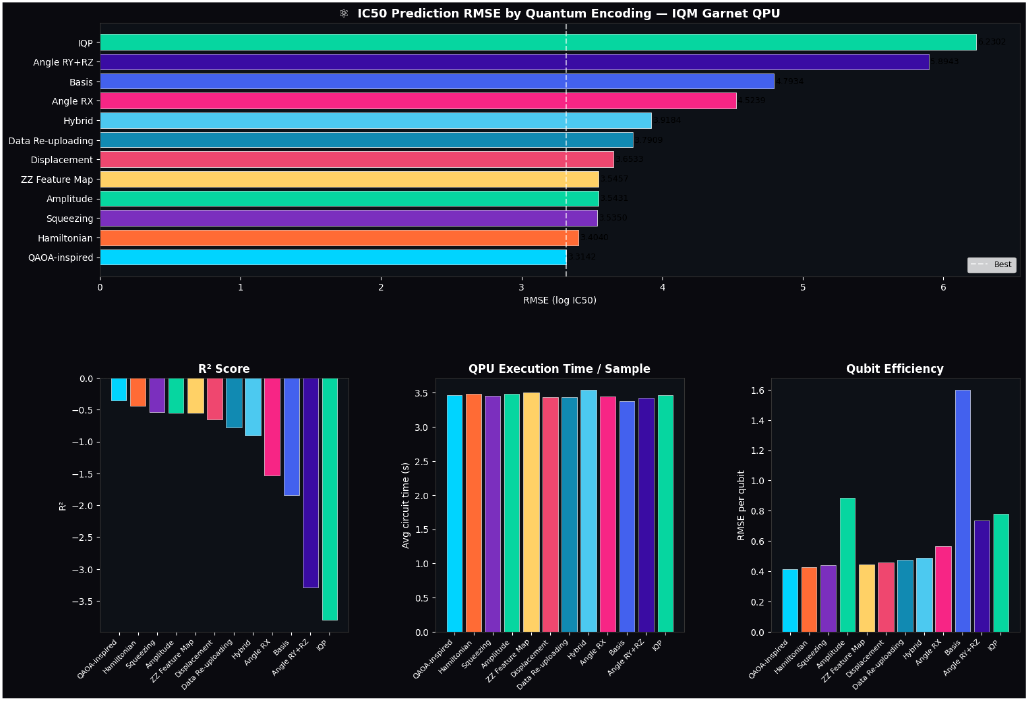
Multi-metric benchmark dashboard. *Top:* RMSE leaderboard (best encoding at bottom); the dashed vertical line marks the QAOA-inspired RMSE of 3.314. *Bottom-left: R*^2^ scores; all are negative, confirming that no encoding surpasses a trivial mean predictor under NISQ conditions. *Bottom-centre:* Average QPU execution time per sample (3.47 ± 0.09 s across all encodings). *Bottom-right:* Qubit efficiency (RMSE/*n*_*Q*_); Amplitude encoding achieves the best ratio (0.886) despite using only 4 qubits.

### D. Statistical Significance

A Wilcoxon signed-rank test was applied to compare the absolute prediction errors of QAOA-inspired (best RMSE) against each of the eleven other encodings. Results are shown in Table III.

**TABLE 3.**
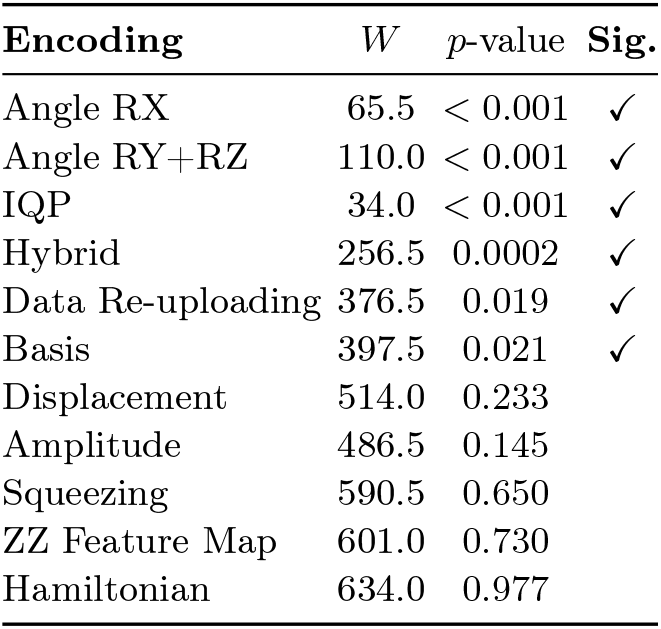
Wilcoxon signed-rank test: QAOA-inspired vs. each competitor. *W* is the test statistic; significance at *α* = 0.05 is indicated.

| Encoding | $W$ | $p$-value | Sig. |
| --- | --- | --- | --- |
| Angle RX | 65.5 | < 0.001 | ✓ |
| Angle RY+RZ | 110.0 | < 0.001 | ✓ |
| IQP | 34.0 | < 0.001 | ✓ |
| Hybrid | 256.5 | 0.0002 | ✓ |
| Data Re-uploading | 376.5 | 0.019 | ✓ |
| Basis | 397.5 | 0.021 | ✓ |
| Displacement | 514.0 | 0.233 |  |
| Amplitude | 486.5 | 0.145 |  |
| Squeezing | 590.5 | 0.650 |  |
| ZZ Feature Map | 601.0 | 0.730 |  |
| Hamiltonian | 634.0 | 0.977 |  |

QAOA-inspired is significantly better than 6 of 11 competitors (*p <* 0.05). Notably, Hamiltonian encoding is the closest competitor (RMSE gap = 0.090, *p* = 0.977): the two are statistically indistinguishable and form a clear performance cluster alongside Squeezing, Amplitude, and ZZ Feature Map (all RMSE *∈* [3.40, 3.55]).

### E. Observed Performance Patterns

a. *Alternating-layer encodings outperform pure-rotation encodings*. QAOA and Hamiltonian encodings, which interleave entanglement with data-dependent phases, achieve RMSE *<* 3.41, versus RMSE *>* 4.52 for pure angle encodings (Angle RX, Angle RY+RZ). We attribute this to the more uniform distribution of data information across the circuit depth, reducing localised noise sensitivity (see also Fig. 3).
b. *Amplitude encoding is qubit-efficient*. Despite using only 4 qubits, Amplitude encoding achieves RMSE = 3.543 and the best qubit-efficiency ratio of 0.886 (Fig. 4, bottom right). This is likely due to the efficient encoding of all 8 features into the 16-amplitude state vector, avoiding the qubit overhead of 8-qubit encodings.
c. *High-expressibility encodings suffer from hardware noise*. IQP and Angle RY+RZ, theoretically the most expressive encodings, rank last by RMSE (6.23 and 5.89, respectively). This is consistent with the noise-induced barren-plateau hypothesis [16]: deep diagonal circuits accumulate more decoherence error on the IQM Garnet’s *T*_1_/*T*_2_ timescales than shallower alternating architectures.
d. *d. All encodings underfit*. The negative *R*^2^ values (Table II) indicate that every encoding performs below a trivial mean predictor. We interpret this as expected behaviour for the NISQ regime at *L* = 2 variational layers and *n* = 50 samples. The benchmark is informative as a *relative* comparison, not as a claim of absolute quantum utility.

## VIII. DISCUSSION

### A. Interpretation in the NISQ Context

All *R*^2^ *<* 0 scores confirm that current NISQ hardware cannot surpass classical baselines for this task at *n* = 50 samples, *L* = 2 layers. This is consistent with the growing consensus that quantum advantage in machine learning requires either fault-tolerant hardware or carefully constructed problem instances with provable kernel advantages [14, 17, 18]. Our results should be read as a *noise-aware hardware benchmark* rather than evidence of quantum utility for drug response prediction.

### B. QAOA-inspired Encoding: Why It Works

The alternating cost–mixer structure provides two complementary mechanisms: (i) the cost layer injects data through *Z*-phase rotations, creating a data-dependent interference pattern; (ii) the mixer layer spreads information globally via uniform *R*_*x*_ rotations. This separation avoids the “concentration of data” problem seen in single-layer angle encodings, where all quantum resources are spent on encoding with none left for mixing.

### C. Limitations

- Regression targets derive from fitting noisy dose-response curves; LN-IC_50_ measurement noise is not accounted for.
- The 8-feature set was chosen for qubit compatibility, not for optimal pharmaceutical prediction.
- COBYLA was limited to 200 evaluations on the classical proxy; a more thorough search may alter rankings.
- No error-mitigation technique (zero-noise extrapolation, readout calibration) was applied.

### D. Recommendations for Practitioners

1. **Prefer alternating-layer encodings** (QAOA-inspired, Hamiltonian) for continuous biomedical regression tasks on superconducting QPUs.
2. **Consider Amplitude encoding** when qubit count is limited; it achieves competitive accuracy with half the circuit width.
3. **Avoid deep diagonal circuits** (IQP) on hardware with limited coherence times. Apply **readout-error mitigation** to improve ⟨*Z*_0_⟩ fidelity before the next hardware campaign.

## IX. RELATED WORK

Havlíček et al. [5] demonstrated kernel-based quantum classification on a 5-qubit IBM device for a synthetic binary task. Schuld and Killoran [4] established the theoretical framework linking feature maps to quantum kernels. Pérez-Salinas et al. [6] introduced data re-uploading as a universal quantum function approximator. The IQP encoding derives from Shepherd and Bremner [11].

For pharmaceutical applications, Batra et al. [19] reviewed QML methods for drug discovery; none involved hardware execution on more than 5 qubits. Recent methodological work by Bowles et al. [22] highlights the subtleties of fairly benchmarking QML models against classical baselines, a concern we address by reporting *R*^2^ relative to a trivial predictor. To our knowledge, the present work is the first hardware benchmark comparing twelve encodings on a biomedical regression task at 8-qubit scale.

## X. CONCLUSION

We have presented a comprehensive, hardware-executed benchmark of twelve quantum encoding strategies for IC_50_ drug-response regression on the IQM Garnet 20-qubit QPU. Our key findings are:

1. **QAOA-inspired encoding** achieves the best RMSE (3.314) and is statistically superior to 6 of 11 competitors (*p <* 0.05).
2. **Alternating cost–mixer architectures** systematically outperform pure-rotation and diagonal encodings under hardware noise.
3. **All encodings under-predict** (*R*^2^ *<* 0), confirming that NISQ-era VQCs require further advances in circuit depth, error mitigation, and training-data scale.
4. The Qrisp/IQM stack provides a reproducible, Python-native pathway for executing variational circuits on superconducting hardware.

Future work will explore (i) readout-error calibration and zero-noise extrapolation, (ii) larger training sets via the IQM mock backend for pre-screening, (iii) transfer learning from classical pre-trained weights, and (iv) extensions to the full GDSC pharmacogenomics feature space.

## ACKNOWLEDGMENTS

The authors thank the IQM Resonance team for providing complimentary QPU credits that made this work possible, and the Qrisp development team for technical support. Cloud QPU access was provided free of charge by IQM Resonance (https://resonance.iqm.tech), including 280 credits awarded following participation in the workshop *Simulating Complex Materials with Quantum Chemistry and SQD*. No external funding was received.

## DATA AVAILABILITY

All code, preprocessed data, raw QPU measurement outputs, trained weights, and figure-generation scripts are publicly available at: https://github.com/rania-derouich/quantum-encoding-ic50-qpu

Detailed circuit definitions, pseudocode, and extended results are provided in the Supplementary Material submitted alongside this manuscript.

GDSC2 dataset: https://www.cancerrxgene.org/downloads/bulk_download

## Notes

### Competing Interest Statement

The authors have declared no competing interest.

## References

[1] J. Biamonte, P. Wittek, N. Pancotti, P. Rebentrost, N. Wiebe, and S. Lloyd, Quantum machine learning, Nature 549, 195 (2017).

[2] M. Schuld and F. Petruccione, Machine Learning with Quantum Computers (Springer, Cham, 2021).

[3] M. Cerezo et al., Variational quantum algorithms, Nat. Rev. Phys. 3, 625 (2021).

[4] M. Schuld and N. Killoran, Quantum machine learning in feature Hilbert spaces, Phys. Rev. Lett. 122, 040504 (2019).

[5] V. Havlíček et al., Supervised learning with quantum-enhanced feature spaces, Nature 567, 209 (2019).

[6] A. Pérez-Salinas, A. Cervera-Lierta, E. Gil-Fuster, and J. I. Latorre, Data re-uploading for a universal quantum classifier, Quantum 4, 226 (2020).

[7] S. Lloyd, M. Schuld, A. Ijaz, J. Izaac, and N. Killoran, Quantum embeddings for machine learning, arXiv:2001.03622 (2020).

[8] N. Killoran et al., Continuous-variable quantum neural networks, Phys. Rev. Res. 1, 033063 (2019).

[9] M. Möttönen, J. J. Vartiainen, V. Bergholm, and M. M. Salomaa, Transformation of quantum states using uniformly controlled rotations, Quantum Inf. Comput. 5, 467 (2005).

[10] M. A. Nielsen and I. L. Chuang, Quantum Computation and Quantum Information, 10th anniversary ed. (Cambridge University Press, 2010).

[11] D. Shepherd and M. J. Bremner, Temporally unstructured quantum computation, Proc. R. Soc. A 465, 1413 (2009).

[12] E. Farhi, J. Goldstone, and S. Gutmann, A quantum approximate optimization algorithm, arXiv:1411.4028 (2014).

[13] K. Mitarai, M. Negoro, M. Kitagawa, and K. Fujii, Quantum circuit learning, Phys. Rev. A 98, 032309 (2018).

[14] M. Schuld and N. Killoran, Is quantum advantage the right goal for quantum machine learning? PRX Quantum 3, 030101 (2022).

[15] F. Iorio et al., A landscape of pharmacogenomic interactions in cancer, Cell 166, 740 (2016).

[16] S. Wang et al., Noise-induced barren plateaus in variational quantum algorithms, Nat. Commun. 12, 6961 (2021).

[17] H.-Y. Huang et al., Power of data in quantum machine learning, Nat. Commun. 12, 2631 (2021).

[18] Y. Liu, S. Arunachalam, and K. Temme, A rigorous and robust quantum speed-up in supervised machine learning, Nat. Phys. 17, 1013 (2021).

[19] K. Batra, K. M. Zorn, D. H. Foil, E. Minerali, V. A. Gawriljuk, T. R. Lane, and S. Ekins, Quantum machine learning algorithms for drug discovery applications, J. Chem. Inf. Model. 61, 2641 (2021).

[20] M. J. D. Powell, A direct search optimization method that models the objective and constraint functions by linear interpolation, in Advances in Optimization and Numerical Analysis, ed. S. Gomez and J.-P. Hennart (Kluwer, Dordrecht, 1994), pp. 51–67.

[21] Qrisp development team, Qrisp: A high-level quantum programming framework, https://qrisp.eu (2024).

[22] J. Bowles, S. Ahmed, and M. Schuld, Better than classical? The subtle art of benchmarking quantum machine learning models, arXiv:2403.07059 (2024).

