## Supplementary Material for "Quantum Encoding Strategies for Drug Response Prediction: An Exhaustive Benchmark on a 20-Qubit Superconducting QPU"

July 8, 2026

**4 Hardware Execution Log Summary** **5**

**5 Additional Figures** **5**

### 1 Circuit Definitions

All circuits are implemented in Qrisp 0.8.2 using `QuantumVariable` objects. The shared variational ansatz appended to every encoding consists of $L = 2$ layers, each comprising $R_y(\theta_{l,i,0})\, R_z(\theta_{l,i,1})$ on every qubit followed by a nearest-neighbour CZ ladder.

#### 1.1 Encoding 1: Angle RX

$$\mathcal{U}_1(x) = \bigotimes_{i=0}^{7} R_x(x_i) \tag{1}$$

Simple single-rotation angle encoding. Low expressibility but hardware-efficient.

#### 1.2 Encoding 2: Angle RY+RZ

$$\mathcal{U}_2(x) = \bigotimes_{i=0}^{7} R_y(x_i)\, R_z(x_{(i+4) \bmod 8}) \tag{2}$$

Two-rotation encoding using complementary feature pairs.

#### 1.3 Encoding 3: Amplitude

Feature vector $x \in \mathbb{R}^8$ is zero-padded and $\ell_2$-normalised to $\tilde{x} \in \mathbb{R}^{16}$, then loaded as quantum state amplitudes on $n = 4$ qubits:

$$\mathcal{U}_3(x)\, |0\rangle^{\otimes 4} = \sum_{k=0}^{15} \tilde{x}_k\, |k\rangle\,. \tag{3}$$

Implemented via `QuantumVariable.qs.init_state()` in the Qrisp statevector simulator, compiled to the QPU backend.

#### 1.4 Encoding 4: Basis

Feature vector is discretised to an integer index $m = \lfloor \sum_i x_i \cdot 2^{7-i} \rfloor \bmod 8$, then loaded as:

$$\mathcal{U}_4(x)\, |0\rangle^{\otimes 3} = |m\rangle_{\text{binary}}\,. \tag{4}$$

Only 3 qubits are used. Minimal depth; no continuous information preserved.

#### 1.5 Encoding 5: ZZ Feature Map

Two Hadamard+phase layers with two-body interaction:

$$\mathcal{U}_5(x) = \left[\prod_i \mathrm{CZ}_{i,i+1}\, e^{i x_i x_{i+1} Z_i} \prod_i P(x_i) \prod_i H_i\right]^2 \tag{5}$$

#### 1.6 Encoding 6: IQP

Instantaneous Quantum Polynomial encoding with diagonal gates:

$$\mathcal{U}_6(x) = H^{\otimes n} \prod_i R_z(x_i) \prod_{\text{even } i} \mathrm{CZ}_{i,i+1}\, R_z(x_i x_{i+1})\, H^{\otimes n} \tag{6}$$

##### 1.7 Encoding 7: Hamiltonian (Trotterised)

Encodes data as Hamiltonian parameters: $H(x) = \sum_i x_i Z_i + \sum_i x_i x_{i+1} Z_i Z_{i+1}$, evolved via 2-step first-order Trotter:

$$\mathcal{U}_7(x) = \left[ e^{-i \sum_i x_i Z_i \frac{t}{2}} \cdot e^{-i \sum_i x_i x_{i+1} Z_i Z_{i+1} \frac{t}{2}} \right]^2 \quad (t = 1) \tag{7}$$

Initial state: uniform superposition $|+\rangle^{\otimes 8}$.

##### 1.8 Encoding 8: QAOA-inspired (Best performer)

Alternating cost and mixer layers:

$$\mathcal{U}_8(x) = \prod_{l=1}^{L} \underbrace{\bigotimes_i R_x(2\beta_l)}_{\text{mixer}} \cdot \underbrace{\text{CZ}_{\text{ladder}} \cdot \bigotimes_i R_z(2x_i)}_{\text{cost layer}} \tag{8}$$

Initial state $|+\rangle^{\otimes 8}$; mixer parameter $\beta_l$ is trainable.

##### 1.9 Encoding 9: Squeezing (DV approximation)

$$\mathcal{U}_9(x) = \text{BS} \cdot \bigotimes_i R_z(x_i^2) \cdot \bigotimes_i R_x(x_i) \tag{9}$$

where BS denotes beam-splitter-like operations: $\text{CZ}_{i,i+1}$, $R_y(\pi/4)_i$, $R_y(-\pi/4)_{i+1}$, $\text{CZ}_{i,i+1}$ for even $i$.

##### 1.10 Encoding 10: Hybrid (Angle + Amplitude)

Qubits 0–3: angle encoding ($R_x(x_i)$). Qubits 4–7: amplitude encoding of $x_{4:8}$, with inter-block CZ bridges.

##### 1.11 Encoding 11: Displacement (CV-DV Bridge)

$$\mathcal{U}_{11}(x) = \bigotimes_i R_y(2\alpha_i \sin\phi_i)\, R_x(2\alpha_i \cos\phi_i), \quad \alpha_i = x_i,\ \phi_i = x_{(i+1) \bmod 8} \tag{10}$$

##### 1.12 Encoding 12: Data Re-uploading

Data $x$ is re-injected at every variational layer $l$:

$$\mathcal{U}_{12}(x) = \prod_{l=0}^{L} \bigotimes_i R_z(x_{(i+l) \bmod 8})\, R_x(x_i) \cdot V_l(\theta_l) \tag{11}$$

#### 2 COBYLA Optimisation Details

**Algorithm.** COBYLA (Constrained Optimisation By Linear Approximations) constructs a simplex of function evaluations and fits a linear model to estimate the gradient direction, without requiring analytical gradients.

**Configuration.**

- Training set: all $n = 50$ samples (classical proxy, no QPU)
- Maximum function evaluations: 200
- Initial trust-region radius: $\rho_0 = 0.5$
- Objective: MSE on training subset
- Initial weights: $\theta_0 \sim \mathcal{U}(-\pi, \pi)$

**Result.** COBYLA reached maximum iterations without meeting the convergence criterion, with final training proxy MSE = 0.0954. This indicates room for improvement with a larger budget or more training samples.

### 3 Dataset Details

#### 3.1 GDSC2 Feature Engineering

The 8 selected features and their preprocessing:

| Feature | Type | Preprocessing |
|---|---|---|
| AUC | Continuous | Min-max $\to [-\pi, \pi]$ |
| RMSE | Continuous | Min-max $\to [-\pi, \pi]$ |
| Z_SCORE | Continuous | Min-max $\to [-\pi, \pi]$ |
| DRUG_ID | Integer | Min-max $\to [-\pi, \pi]$ |
| COSMIC_ID | Integer | Min-max $\to [-\pi, \pi]$ |
| cancer_UNCLASSIFIED | Binary | $\{-\pi, \pi\}$ |
| cancer_LUAD | Binary | $\{-\pi, \pi\}$ |
| cancer_SCLC | Binary | $\{-\pi, \pi\}$ |

All encodings use $[-\pi, \pi]$ via the `QuantumPreprocessor` default. The amplitude encoding normalises the feature vector to unit $\ell_2$ norm.

#### 3.2 Stratified Sampling

50 samples were drawn using percentile-stratified sampling over LN-$IC_{50}$ with 10 equally-spaced percentile bins (5 samples per bin), ensuring coverage of the full response range in the QPU subset.

### 4 Hardware Execution Log Summary

| Encoding | Total time (s) | Avg/sample (s) |
|---|---|---|
| Angle RX | 172.7 | 3.45 |
| Angle RY+RZ | 171.5 | 3.43 |
| Amplitude | 173.9 | 3.48 |
| Basis | 169.1 | 3.38 |
| ZZ Feature Map | 175.1 | 3.50 |
| IQP | 173.5 | 3.47 |
| Hamiltonian | 174.0 | 3.48 |
| QAOA-inspired | 173.3 | 3.47 |
| Squeezing | 173.0 | 3.46 |
| Hybrid | 176.8 | 3.54 |
| Displacement | 172.2 | 3.44 |
| Data Re-uploading | 171.9 | 3.44 |
| **Total** | **2076.0** | **3.46 (mean)** |

Total QPU wall time: $\approx$ 34.6 minutes for all 600 circuit submissions (12 encodings $\times$ 50 samples).

### 5 Additional Figures

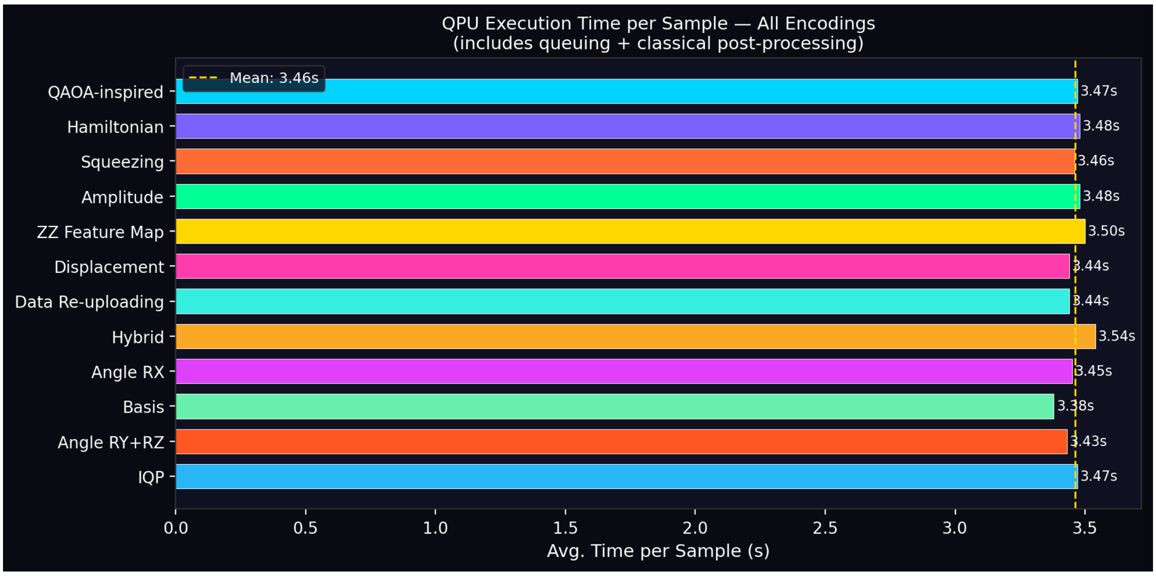

Figure 1: QPU execution time per sample for each encoding. Mean latency: $3.47 \pm 0.09$ s.

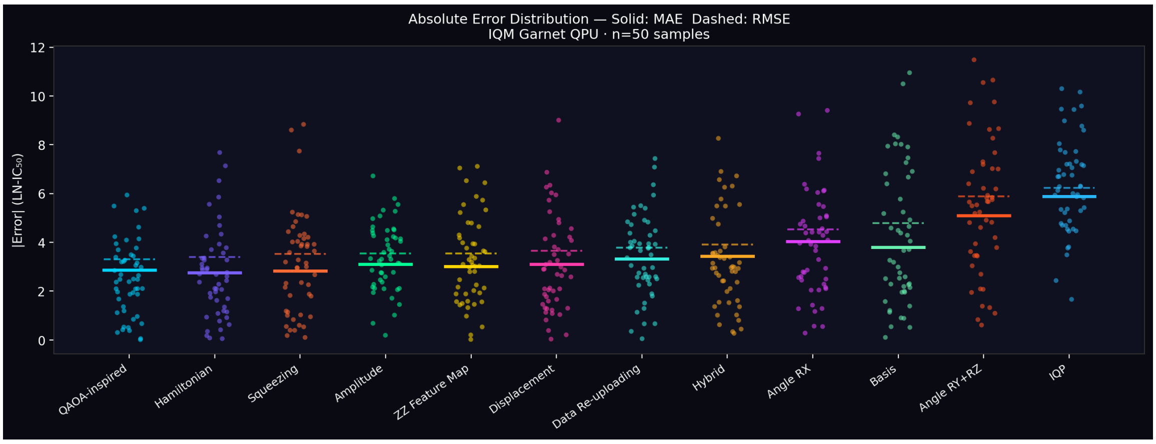

Figure 2: Absolute error distribution per encoding. Solid lines: MAE; dashed lines: RMSE. Points are individual sample errors (with jitter).

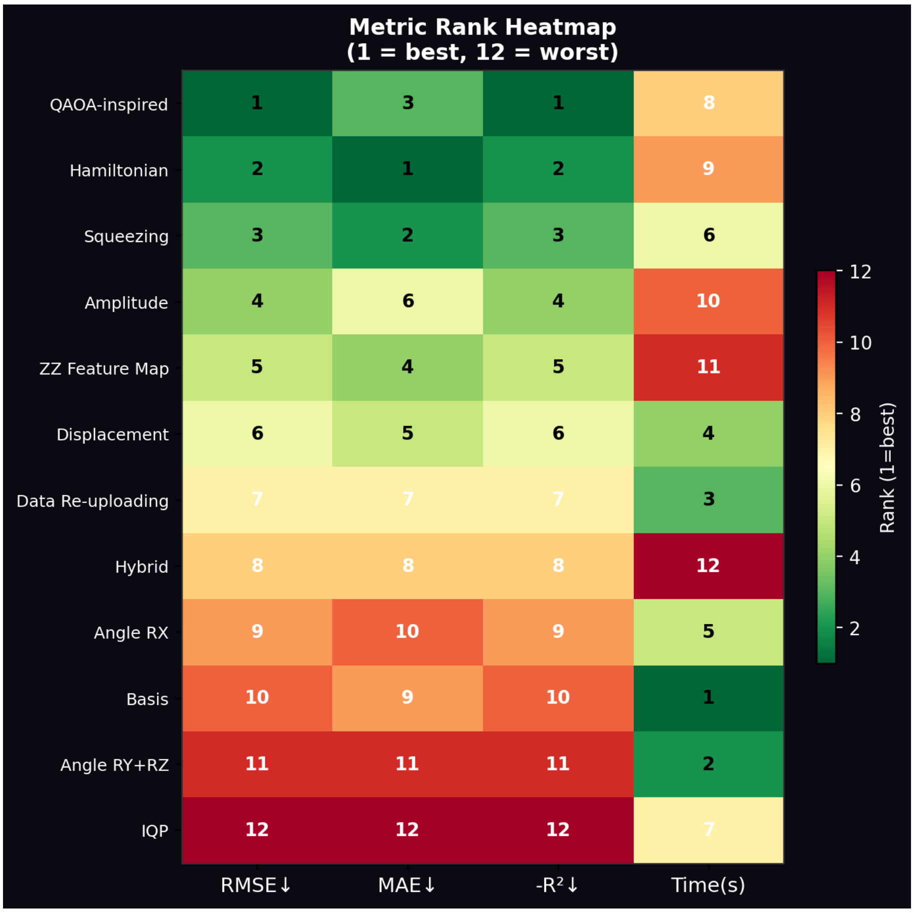

Figure 3: Metric rank heatmap (1 = best, 12 = worst) across RMSE, MAE, $-R^2$ and execution time. QAOA-inspired ranks first on RMSE and $-R^2$; Hamiltonian ranks first on MAE; Basis ranks first on execution time.
